# An automated approach to improve the speed and accuracy of pericyte and microglia quantification in whole mouse brain sections

**DOI:** 10.1101/2021.04.29.442048

**Authors:** Jo-Maree Courtney, Gary P Morris, Elise M Cleary, David W Howells, Brad A Sutherland

**Author notes:** **Author Contributions:** JMC, GPM, DWH, and BAS designed research; JMC, GPM and EMC performed research; JMC and GPM analysed data; JMC, GPM and BAS wrote the paper. **Correspondence:** Brad A. Sutherland.

## Abstract

Whole slide scanning technology has enabled the generation of high-resolution images of complete tissue sections. However, commonly used analysis software is often unable to handle the large data files produced. Here we present a method using the open-source software QuPath to detect, classify and quantify fluorescently-labelled cells (microglia and pericytes) in whole coronal brain tissue sections. Whole brain sections from both male and female NG2DsRed x CX_3_CR1^+/GFP^ mice were analysed. Small regions of interest were selected and manual counts were compared to counts generated from an automated approach, across a range of detection parameters. The optimal parameters for detecting cells and classifying them as microglia or pericytes in each brain region were determined and applied to annotations corresponding to the entire cortex, hippocampus, thalamus and hypothalamus in each section. 3.71% of all detected cells were classified as pericytes, however this proportion was significantly higher in the thalamus (6.39%) than in other regions. In contrast, microglia (4.45% of total cells) were more abundant in the cortex (5.54%). No differences were detected between male and female mice. In conclusion, QuPath offers a user-friendly, rapid and accurate solution to whole-slide image analysis which could lead to important new discoveries in both health and disease.

**Significance Statement:** Rapid and accurate quantification of cell numbers and distributions from whole tissue sections represents a difficult challenge in biomedical research. Slide scanning microscopes generate high-resolution images of complete tissue sections but most common image analysis software packages struggle to cope with the large data files they produce. We provide a method for rapidly and accurately quantifying pericyte and microglia cell numbers in whole brain tissue sections using QuPath, a new open-source software designed specifically to overcome this challenging roadblock.

## Introduction

The mammalian brain is a large and complex organ with numerous cell types. The parenchymal cells of the brain, including neurons, microglia, astrocytes, oligodendrocytes, and oligodendrocyte precursor cells (OPCs), co-exist alongside cells lining the walls of the ventricles (ependymal cells), and cells forming the blood vessels of the brain (endothelial cells, pericytes, and vascular smooth muscle cells). Accurately determining the density and spatial relationships between these different cell types, in any given brain region, can provide clues to the importance and functions of each cell type in both health and disease.

Brain cells can be visualised by many forms of microscopy including brightfield or fluorescence microscopy following histological or immunohistochemical processing of isolated cells or tissue sections. The quantitative and spatial analysis of cells has traditionally been limited by the field of view of the microscope and the workload associated with analysis of multiple fields of view, which can hinder the detection of patterns across larger regions. Virtual microscopy technology, which enables the scanning of whole microscope slides at high resolution, has emerged in the last two decades to overcome this limitation (Al-Janabi et al., 2012). Although whole slide scanning has predominantly been adopted in the field of diagnostic pathology, basic research laboratories can also benefit from the analysis of different cell types over large areas of tissue.

Whole slide scanning has however created a new roadblock. The files generated by slide scanning microscopes are large and difficult to handle for many common image analysis programs, including open-source software such as ImageJ and CellProfiler. To overcome this, researchers resort to reducing image resolution (which can reduce the accuracy of the analysis), sampling smaller regions of interest (as a representative of the larger whole), or painstakingly analysing the whole by consecutively imaging and analysing small regions of interest. Recently, the open-source application QuPath was specifically developed to better enable pathologists and researchers to analyse whole slide images (Bankhead et al., 2017). QuPath can integrate with ImageJ and other packages to reuse carefully developed analysis tools and allows the user to rapidly analyse high resolution images without requiring expensive, specialised computing facilities and without having to rely on sampling smaller regions of interest. The initial application for QuPath was for tumor identification and biomarker evaluation in cancer (Bankhead et al., 2018; Humphries et al., 2018; Ledys et al., 2018; Loughrey et al., 2018), but its extensible platform provides the flexibility to analyse large and complex images across a range of biomedical settings (Bankhead et al., 2017). For example, the effective identification of GFAP-positive astrocytes across whole brain sections recently provided the first demonstration of the use of QuPath in neurological microanatomy (Finney et al., 2020). Here, we demonstrate the potential of QuPath to rapidly and accurately detect fluorescently-labelled brain cells, in particular microglia and pericytes.

Both microglia and pericytes are distributed widely throughout the brain and have important functions in health and disease. Microglia, historically considered the innate immune cells of the brain (Morris et al., 2013), are unique to the central nervous system (CNS), with key roles in sculpting, maintaining and modifying neural circuitry through their influence on synaptic and structural plasticity (Colonna and Butovsky, 2017). Pericytes, a cell present throughout the central nervous system and the periphery, have numerous roles in the brain including the regulation of cerebral blood flow (Hall et al., 2014) and the maintenance of the blood-brain barrier (Armulik et al., 2010). The historical and contemporary research on both cell types in health and disease have been extensively reviewed elsewhere (Beard et al., 2020; Brown et al., 2019; Colonna and Butovsky, 2017; Morris et al., 2013; Sweeney et al., 2016).

In this study, we have used QuPath to quantify the relative numbers of microglia and pericytes in whole coronal brain tissue sections derived from transgenic mice expressing fluorescently-labelled microglia and pericytes. We describe, for the first time, the use of QuPath to analyse images of whole mouse brain sectionss for fluorescently-labelled cells in an automated fashion. We also highlight optimisation processes that permit quantification of microglia and pericytes under different imaging circumstances in different brain regions. Our approach can be easily applied to any brain region or other tissue types, or other fluorescently-labelled cells, and can be used to quantify cell numbers in different disease states.

## Materials and Methods

### Animals, tissue acquisition and processing

All animal procedures were approved by the Animal Ethics Committee, University of Tasmania (A0018608) and conformed with the Australian NHMRC Code of Practice for the Care and Use of Animals for Scientific Purposes –2013 (8th Edition). Hemizygote NG2DsRed transgenic mice (Jackson Laboratories Stock #008241) were backcrossed onto a C57BL/6J background and crossbred with CX_3_CR1^GFP/GFP^ transgenic mice (Jackson Laboratories Stock #005582, C57BL/6J background) to produce NG2DsRed x CX_3_CR1^+/GFP^ mice. Mice were group housed in Optimouse caging on a 12h light:dark cycle (lights on: 0700-1900) with *ad libitum* access to standard chow and water.

Eight 12-week-old male and female NG2DsRed x CX_3_CR1^+/GFP^ mice weighing 18.7-30.3 g (Table 1) were killed with a lethal intraperitoneal injection of pentobarbitone (300 mg/kg) and immediately transcardially perfused with 4% paraformaldehyde (PFA, pH 7.4). Whole brains were harvested and additionally post-fixed in 4% PFA for 1.5h, then transferred to 30% sucrose in 1x PBS until they sank. Whole brains were embedded in Cryomatrix embedding resin (Shandon, Cat# 6769006) and frozen at −80°C until cryosectioning. 40μm coronal sections were cut at −18°C using a cryostat and placed free floating in 1x PBS. Using the Allen Brain Atlas as a guide (Lein et al., 2007), tissue sections −1.70 mm from Bregma were mounted onto microscope slides (Dako, Cat# K802021-2), allowed to dry upright for 30 min, washed for 5 min in 1x PBS Tween (0.1%), rinsed for 30s in 1x PBS followed by 10s in distilled H_2_O, air dried for 5 min, then coverslipped with Prolong Gold antifade reagent with DAPI (Life Technologies, Cat# P36935).

**Table 1.** Descriptive statistics of animals and tissue. All statistics are mean ± standard deviation. Male and Female groups were compared with an unpaired t-test. * p < 0.05, *** p < 0.0005, ns = not significant.

|  | Male (n=3) | Female (n=5) | P-value |
| --- | --- | --- | --- |
| **Weight (g)** | 27.77 ± 2.37 | 19.74 ± 1.05 | p = 0.0005 *** |
| **Age (days)** | 87.67 ± 2.52 | 85.00 ± 0.00 | p = 0.0457 * |
| **Tissue slice area (mm<sup>2</sup>)** | 42.17 ± 0.86 | 42.03 ± 1.25 | p = 0.8739 ns |

### Image acquisition

Images were acquired using a VS120 Virtual Slide System (Olympus). Whole slides were first scanned in the DAPI channel (Ex: 388nm; Em: 448nm) at 2x magnification and then the outlines of whole coronal tissue sections were traced for scanning at 40x magnification. A focus-map (the highest density possible) was auto-generated across the entirety of each coronal section and the plane of focus was automatically determined based on the DAPI channel. DAPI (Ex: 388nm; Em: 448nm), DsRed (Ex: 576nm; Em: 625nm) and GFP (Ex: 494nm; Em: 530nm) signals were imaged in the same focal plane. Optimum exposure times were initially determined manually and then kept consistent for all images (DAPI: 50 ms, DsRed: 100 ms, GFP: 50ms). The .vsi files generated were approximately 2GB each in size.

### Image Analysis – computing and software

All image analysis was performed on a standard desktop computer with an Intel Core i7-6700 processor and 16GB installed memory running Windows 10 and QuPath-0.2.3. Script development was aided by IntelliJ IDEA 2020.3.2 (Community edition). Analysis of exported data was performed using Microsoft Excel and GraphPad Prism 9.0.2.

The method described here was based on the Multiplexed Analysis Tutorial found in the QuPath Online Documentation (Bankhead, 2020) with adaptations made for whole brain section analysis and the specifics of the tissue used here. An overview of the analysis pipeline is shown in Figure 1.

**Figure 1.**
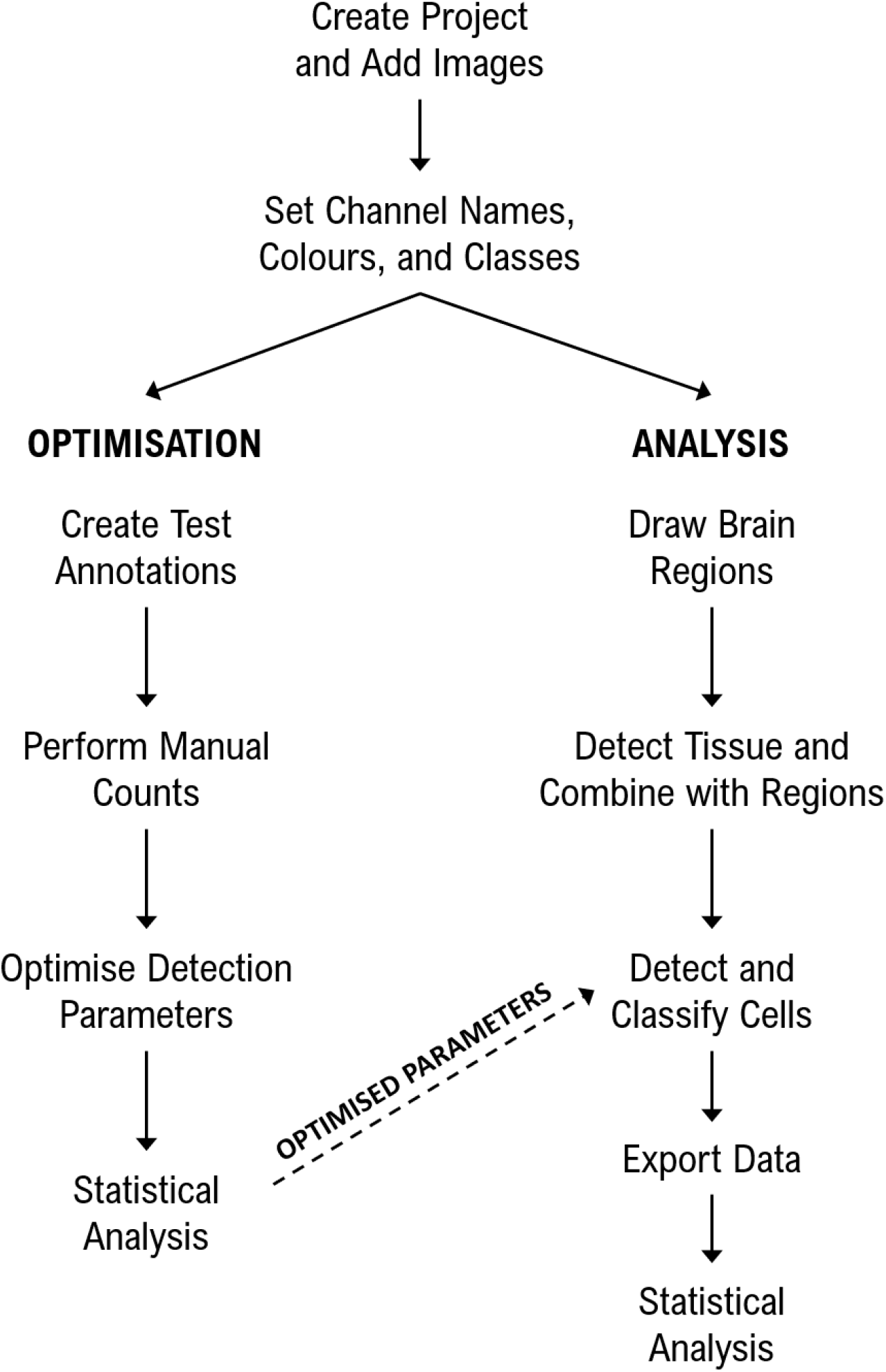
Analysis Pipeline. Flowchart summarising the steps taken to optimise analysis parameters and then to detect and classify fluorescently labelled cells in whole mouse brain sections in QuPath.

### Code Accessibility

Source code for the scripts and classifiers used here are available upon request.

### Image analysis –project setup

A QuPath project was created to allow the application of scripts and classifiers across multiple images. All .vsi files were loaded with the image type set to ‘fluorescence’. QuPath does not hold the actual image files but rather links to the original images, and so it was ensured that the project file and original image files were never separated. The project was duplicated to create separate projects for optimisation and post-optimisation analysis.

### Image analysis – channel names, colours, and classes

Appropriate channel colours and names (DAPI, DsRed, GFP) were set for all images as a batch using the script ‘Channels and Colours.groovy’ and classes were created from these channel names using the ‘Populate from Image Channels’ command (Figure 2A). In our figures the DAPI, DsRed and GFP have been pseudo coloured blue, magenta and green, respectively.

**Figure 2.**
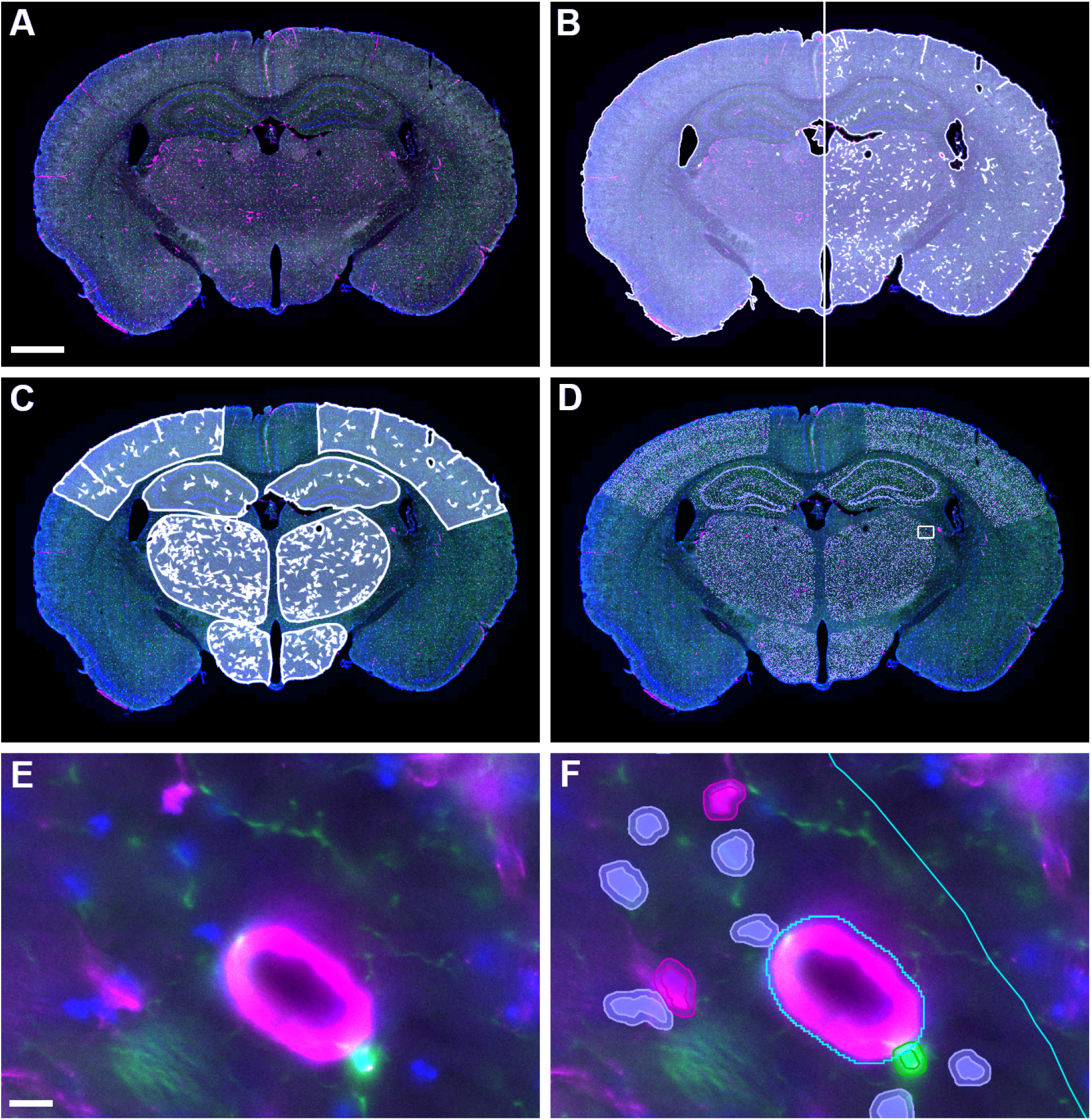
Stages of QuPath analysis on whole mouse brain sections. **A)** Imported image following correction of channel colours; **B)** Initial tissue detection (left) and following subtraction of large vessels and edges (right); **C)** Brain regions intersected with detected tissue; **D)** Overlay of detected nuclei (grey) on tissue; **E)** High magnification view of a region of the thalamus (indicated by box in D); **F)** with annotation boundaries in cyan and detected nuclei outlined in magenta (Ds-Red), green (GFP) and grey (other). Scale bars represent 5mm (A-D) and 10μm (E-F). DAPI, GFP and DsRed signal are coloured blue, green and magenta respectively. Extended Data Figure 2-1 shows an example of the placing of annotations for manual cell counting.

### Optimisation – Selection of test annotations and manual counting

QuPath has the functionality to analyse entire brain sections or smaller regions of interest, and different regions of the brain may require different parameters for optimal cell detection. To determine the optimal parameters for our DAPI-positive nuclei detection and the GFP-positive/DsRed-positive cell classifications across the brain, for each image (n=8) a small annotation (300 x 200μm) was drawn in each of six brain regions of interest: upper cortex (layers 1-3), lower cortex (layers 4-6), hippocampus (including dentate gyrus), hippocampus (including CA1/CA3 boundary), thalamus, and hypothalamus (for an example, see Figure 2-1 in Extended Data). In each annotation area, the total number of cells (DAPI-stained nuclei) and the number of DsRed-positive and GFP-positive cells were counted manually by an experienced researcher (GPM) using the points annotation function of QuPath.

### Optimisation of cell detection parameters

In QuPath, fluorescent cell detection can be performed using any channel but most commonly utilises a nuclear stain such as DAPI to first detect all cells. The in-built Cell Detection algorithm requires the selection of various parameters. It is possible to rigorously optimise each of these parameters (i.e. pixel size, background radius, median radius, sigma, min area, max area and threshold) individually, or by mixing and matching different settings for each, then comparing these settings to manual counts to ensure accurate automated detection of cells. Manually changing these parameters to test each possible combination is time consuming, so we designed a script capable of doing this automatically (‘Optimisation of cell detection.groovy’).

To illustrate the importance of optimising cell detection parameters we present our optimisation of one key detection parameter: the DAPI intensity threshold. For simplicity, the other cell detection parameters were kept to QuPath’s defaults, except for Sigma =1.5 and Cell Expansion = 2μm (a cell expansion allows for the detection of fluorescent labelling outside of the nucleus).

For each test annotation, DAPI intensity thresholds were tested in increments of 25, beginning at 50 and ending at 200. The number of detected cells at each threshold was compared to manual counts of DAPI-positive cells to determine the optimal threshold for each region of the brain.

### Optimisation of fluorescent intensity thresholds for cell detection

After optimising the cell detection parameters, we optimised the intensity threshold parameters for classifying detected cells as DsRed-positive or GFP-positive. The inbuilt Positive Cell Detection plugin was applied to the same small annotations used for optimising DAPI cell detection. Here the DAPI threshold for nuclear detection was set to the previously determined optimum for each specific brain region while the threshold (measuring the mean value in the cell) was tested using a script (‘Optimisation of cell classification.groovy’): first for DsRed (thresholds between 200 and 450 in increments of 25) and then for GFP (thresholds between 100 and 300 in increments of 25).

As with DAPI detection, the number of DsRed- and GFP-positive cells detected at each threshold was compared to the manual counts (with visual verification) to determine the optimal DsRed and GFP thresholds for each brain region.

### Image Analysis – Annotation of Brain Regions

For each image in the analysis project, the brush tool was used to draw annotations for the cortex, hippocampus, thalamus and hypothalamus in both left and right hemispheres using the Allen Mouse Brain Atlas as a guide (Lein et al., 2007). The selected regions in the left and right hemispheres were merged to form a single annotation for each region and each annotation was named appropriately using the ‘Set Properties’ dialog box. Two scripts were used to assist this process: ‘Save Annotations.groovy’ exports the annotations for the first image to a file which can then be imported back into the remaining images with ‘Import Annotations.groovy’. Annotations for each image were individually adjusted using the brush tool to fit the specific anatomy of each section.

### Image Analysis – Tissue and Vessel Detection

For each tissue section, the tissue area was defined using a pixel classifier based on the average value of the three channels at high resolution with a Gaussian prefilter, smoothing sigma = 2.0 and threshold = 50 and an annotation created with a minimum area of 1,000,000μm^2^ and a minimum hole size of 1,000μm^2^. This annotation was eroded by 40μm (using ‘Expand Annotations’ set to −40μm) to reduce the effects of tissue processing artefacts around the edge of the tissue, and fragments less than 10,000μm^2^ were removed (using ‘Remove fragments and holes’). In order to exclude large DsRed-positive vessels, likely reflecting DsRed-positive vascular smooth muscle cells, rather than capillary pericytes, a second pixel classifier was created using the DsRed channel at high resolution with a Gaussian prefilter, smoothing sigma = 2.0 and threshold = 400. An annotation was created from this classifier with a minimum size of 150μm^2^ and a minimum hole size of 1,000μm^2^. Next, the Vessels annotation was subtracted from the Tissue annotation. A script was created to incorporate the pixel classifiers and automate these steps (‘Tissue Detection.groovy’) and run as a batch for the project (Figure 2B). Finally, the intersection between the detected tissue (minus large vessels) and the pre-defined brain region annotations was calculated using the script ‘Intersect ROIs.groovy’ (Figure 2C).

### Image Analysis – Detection and Classification of Cells

To accommodate the need for different colour thresholds in different brain regions, a specific composite object classifier was created for each brain region and saved in the QuPath project’s ‘object_classifiers’ folder.

Finally, cell detection and classification was combined into a single script to run for the whole project (‘Cell Detection and Classification.groovy’), using the pre-determined DAPI, DsRed and GFP thresholds for each region (Figure 2D). Examples of cell detections are shown in Figure 2E, F and Figure 3.

**Figure 3.**
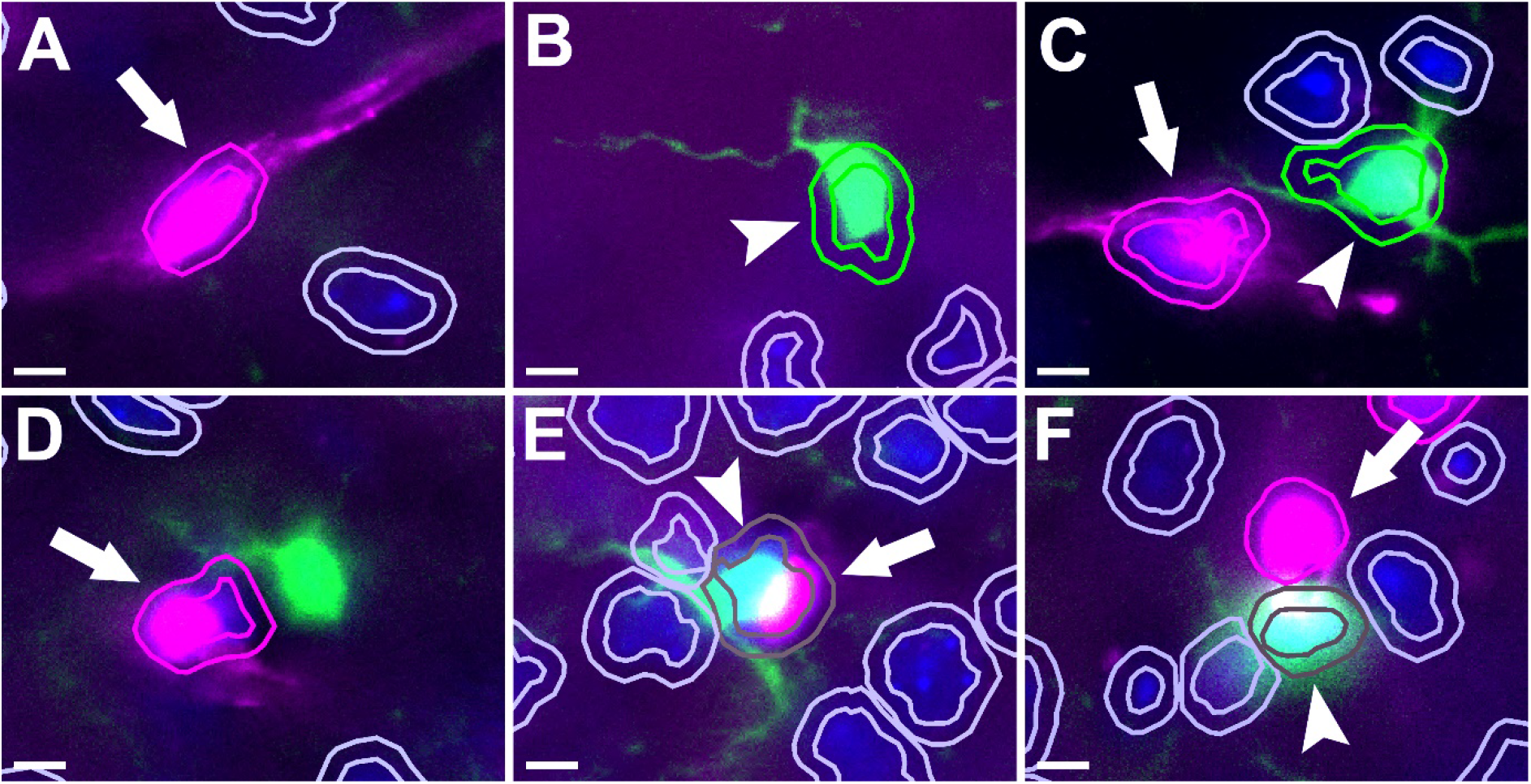
Examples of detected cells. DsRed-positive pericytes are indicated with arrows, GFP-positive microglia with arrowheads. Each cell detected using DAPI staining is shown as an inner ring (nucleus) and outer ring (2μm expansion) coloured according to classification (magenta = DsRed-positive, green = GFP-positive, brown = DsRed- and GFP-positive, grey = DsRed- and GFP-negative). **A-C)** Appropriately classified cells. **D)** The pericyte is classified appropriately but the microglia is not detected due to the nucleus being out of the plane of the section. **E)** A microglia and pericyte that are in close contact and were not able to be separated by the nuclear detection leading to a dual classification. **F)** The pericyte is appropriately classified but enough of the DsRed fluorescence has colocalised with the microglia to cause a dual classification. This figure illustrates two of the possible reasons for cells to be dual-classified, however overall occurrence of dual-classified cells is low (see Figure 5B). Scale bars = 5μm.

### Image Analysis – Export of Measurements

Annotation measurements including area and number of detections were exported for each brain region using QuPath’s Measurement Exporter.

### Statistical analysis

Statistical analysis was performed using GraphPad Prism 9.0.2. Animal and tissue slice measurements were compared with unpaired t-tests. Manual counts were compared to automated counts using the Pearson’s correlation coefficient. Cell counts were compared between brain regions by repeated measures one-way ANOVA with post-hoc Tukey’s multiple comparison tests. Sex effects were tested with repeated measures two-way ANOVA. A p < 0.05 was considered statistically significant.

## Results

### Optimisation

After running the QuPath Cell Detection algorithm through our custom script, we compared manual cell counts from 300 x 200 μm annotations in six different regions to the number of DAPI-positive nuclei detected by the algorithm across seven different DAPI intensity thresholds (summarised in Figure 4A, see extended data Figure 4-1 for individual comparisons). Optimal DAPI-thresholds were selected as the thresholds which provided the greatest accuracy (i.e. closest mean to manual counts) and least variability (i.e. smallest standard deviation), with a preference for undercounting (false negatives) rather than overcounting (false positives). For the cortex, thalamus and hypothalamus, the optimal threshold was determined to be 150, while the hippocampal regions required lower thresholds between 50-100 for accurate cell detection. For the hippocampus, we ideally required a threshold that would be applicable to the whole region (i.e. including CA1/CA3 and DG in the same analysis). We therefore chose a threshold of 75 for the entire hippocampus, which provided accurate counts in both regions. For each brain region, the number of cells detected using the optimised DAPI thresholds significantly correlated to number of cells counted manually (Figure 4D and extended data Figure 4-2).

**Figure 4.**
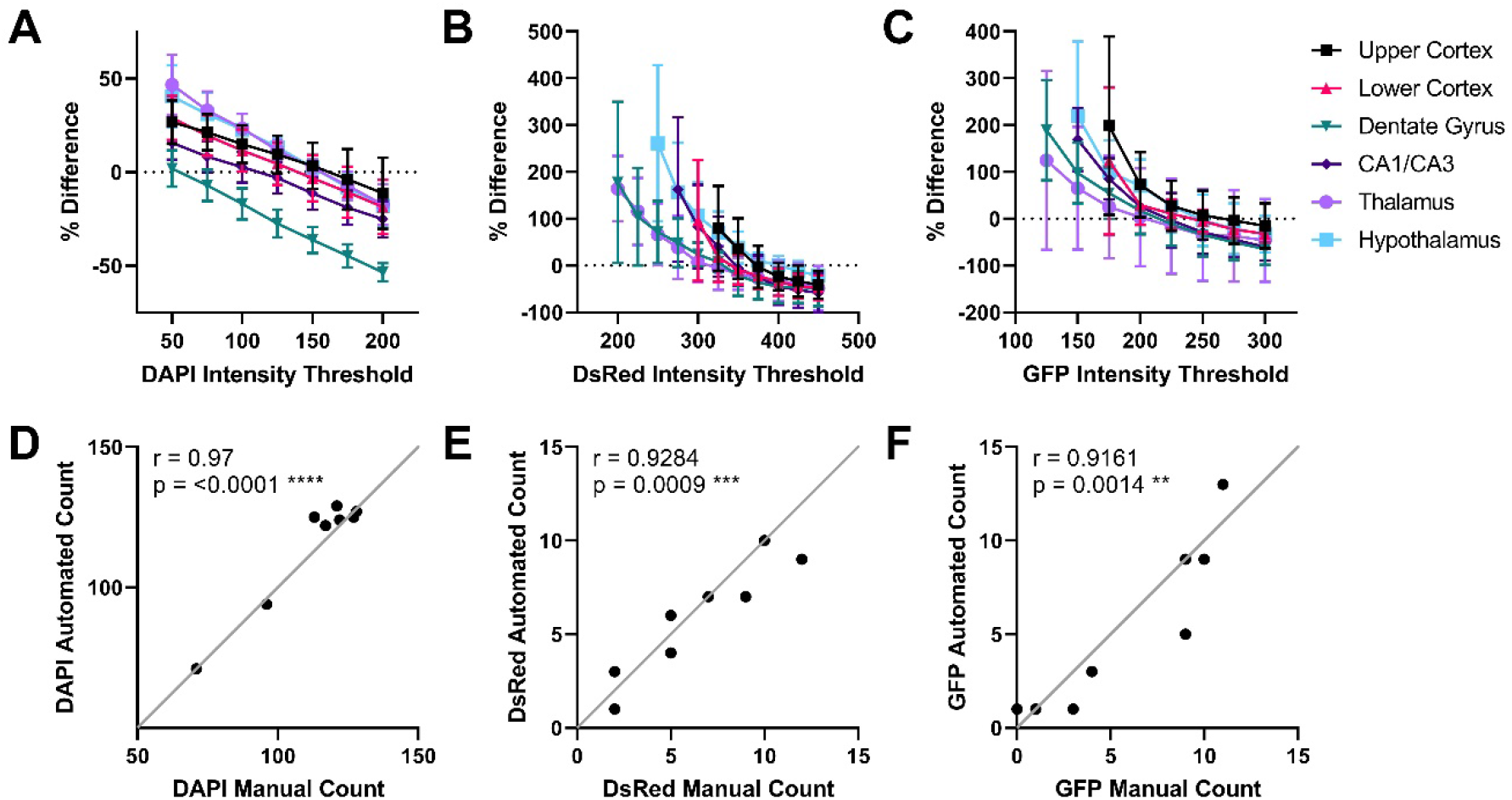
Optimisation of cell detection and classification thresholds. Counts generated by QuPath’s Cell Detection/Positive Cell Detection algorithm were compared to manual cell counts to generate a % difference (dotted line at 0%) with a range of intensity thresholds across six brain regions for **A)** DAPI, **B)** DsRed, **C)** GFP (n=8, mean ± standard deviation). Data for thresholds with standard deviations over 200 have been excluded for the sake of clarity. Extended Data Figure 4-1 shows intensity threshold analyses for annotations of individual brain regions. Example correlations of automated counts to manual counts for the thalamus using the final optimised values for **D)** DAPI, **E)** DsRed, and **F)** GFP. Extended Data Figure 4-2 includes correlation charts for all optimised brain regions.

Next, we used another custom script to iteratively apply QuPath’s Positive Cell Detection algorithm, using the DAPI thresholds optimised for each brain region, to test multiple DsRed and GFP intensity thresholds. As with the optimisation of DAPI thresholds, we compared these to manual counts to determine the optimal thresholds for DsRed- and GFP-positive cell detection in each region (summarised in Figure 4B-C, see extended data Figure 4-1 for individual comparisons). Again, thresholds were selected for accuracy, low variability, and with a preference for false negatives. The number of cells detected using the optimised DsRed and GFP thresholds correlated to number of DsRed- and GFP-positive cells counted manually (Figure 4E-F and extended data Figure 4-2).

The final optimised thresholds for each channel and region are listed in Table 2. Note, these optimised thresholds are only applicable to our specific tissue and would be expected to vary in each laboratory based on the tissue processing methodology and imaging parameters.

**Table 2.** Optimised intensity thresholds for cell detection and classification by brain region.

|  | DAPI | DsRed | GFP |
| --- | --- | --- | --- |
| Cortex | 150 | 375 | 250 |
| Hippocampus | 75 | 350 | 225 |
| Thalamus | 150 | 325 | 200 |
| Hypothalamus | 150 | 400 | 250 |

### Cell Detection and Quantification in Different Brain Regions

Following the optimisation of cell detection parameters on small annotations, we applied our cell detection script with specific Object Classifiers to each region of interest within our coronal sections (1 section per animal, n=8). Automated cell detection (based on DAPI staining) was performed on each region of interest in each brain section (Figure 5A). The total number of cells per area differs significantly between brain regions (one-way ANOVA, *p* = 0.004).

**Figure 5.**
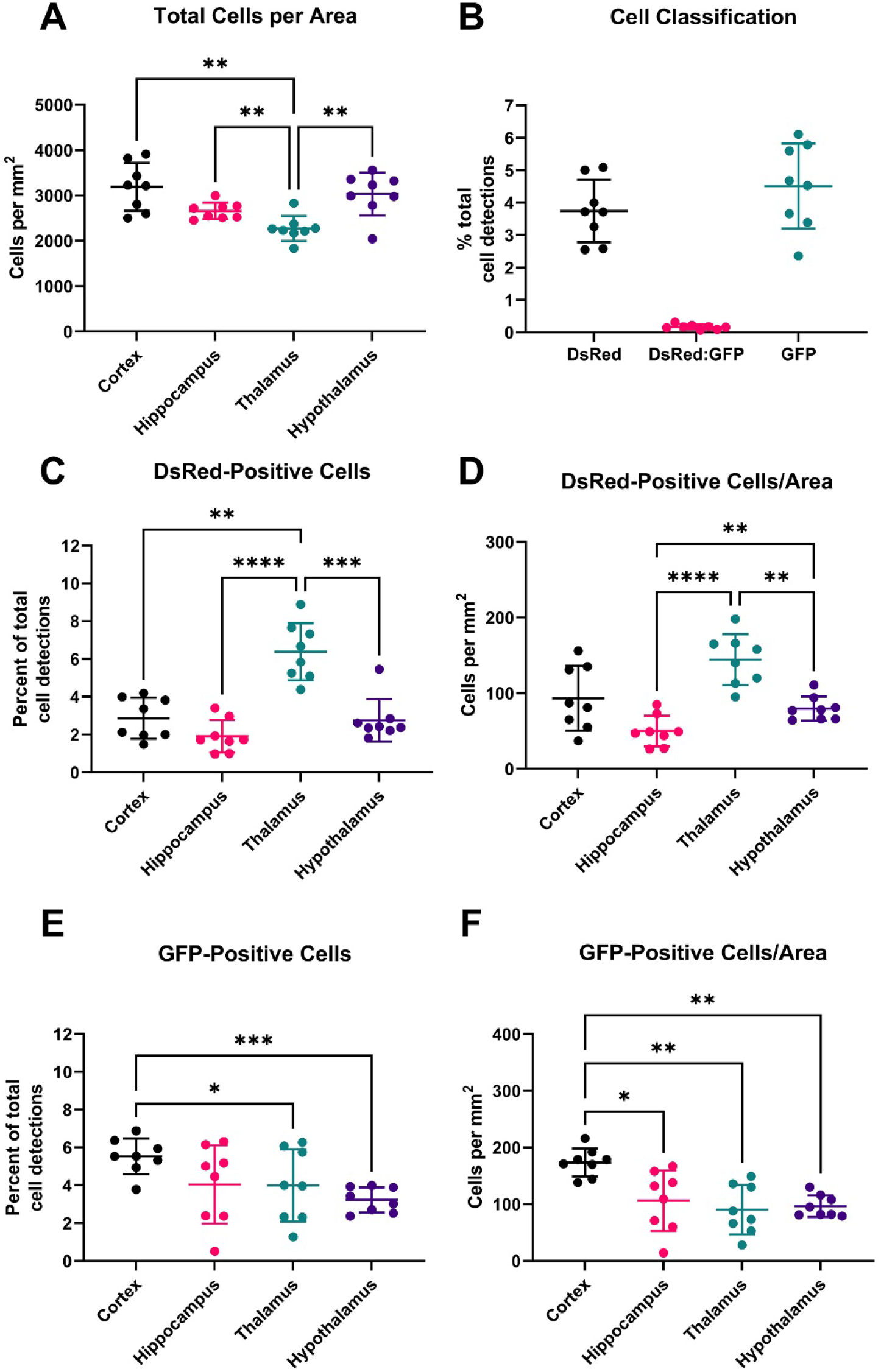
Detection and classification of cells. **A)** Total cells detected per mm^2^ tissue area in each brain region by DAPI nuclei staining. **B)** Percentage of cells by classification across all brain regions measured. Ds-Red positive cells by brain region **C)** as percentage of total cell detections and **D)** per mm^2^. GFP positive cells by brain region **E)** as percentage of total cell detections and **F)** per mm^2^. Statistical analysis by repeated measures one-way ANOVA with post-hoc Tukey’s multiple comparison test. * p < 0.05, ** p < 0.01, *** p < 0.001. Extended Data Figure 5-1 shows detection and classification analyses comparing male and female mice with no statistically significant differences observed.

Across all regions tested, 3.74% (±0.90%) detected cells were classified as DsRed-positive (pericytes), 4.51% (±1.23%) were classified as GFP-positive (microglia), and 0.17% (±0.07%) were classed as positive for both DsRed and GFP (Figure 5B). One reason for the classification of a subset of cells as positive for both markers is the close proximity of some pericytes to microglia, making it difficult for the automated analysis to distinguish individual cells that have overlapping DsRed and GFP fluorescence (see Figure 3E,F for examples). Given the small numbers, these cells were excluded from further analysis.

The proportion of total DAPI-positive cells that were identified as DsRed- or GFP-positive differed significantly between brain regions. The thalamus had a significantly higher proportion of DsRed-positive pericytes (6.39% ± 1.51%) compared to the other brain regions assessed (cortex: 2.86% ± 1.08%, p = 0.0017; hippocampus: 1.91% ± 0.86%, p <0.0001; and hypothalamus: 2.76% ± 1.13%, p = 0.0007) (Figure 5C). The cortex had a significantly higher proportion of GFP-positive microglia (5.54% ± 0.94%) than other brain regions assessed (hippocampus: 4.05% ± 2.07%, p = 0.0597; thalamus: 3.99% ± 1.91%, p = 0.0311; and hypothalamus: 3.23% ± 0.66%, p = 0.0001) (Figure 5E). A similar pattern of regional differences was evident when cells counts were expressed as cells/mm^2^ of tissue area (Figure 5 D,F).

No statistical differences were identified between male and female mice for any of the cell detection or classification measures described (Extended Data Figure 5-1).

## Discussion

In this study, we have detailed a fully automated method to identify fluorescently labelled nuclei, microglia and pericytes in high-resolution images using QuPath. To validate our approach we compared automated nuclei, pericyte and microglia counts to manual counts. The approach we described offers an unbiased, replicable method to quantitate nuclei and cell numbers across large fluorescently labelled tissue sections, drastically reducing the time taken to obtain cell counts. Below we discuss the importance of optimising the QuPath cell detection parameters for each project. Furthermore, we highlight limitations in our QuPath methodology and we discuss other useful features of QuPath beyond those we have assessed in this work.

QuPath represents a significant advance in biomedical image interpretation by enabling batch analysis of large (> 2GB), pyramidal image files produced by slide scanners in a scriptable, open-source environment on a standard desktop computer without the need to downsample or limit analysis to small regions of interest. Since its release in 2017, the vast majority of work published using QuPath has analysed tissue sections stained with chromogenic immunohistochemistry. Here we provide the first investigation utilising QuPath to detect fluorescently labelled brain cells in mouse brain tissue sections, specifically pericytes expressing DsRed and microglia expressing GFP.

When using automated approaches for cell detection and quantification it is prudent to optimise the automated cell detection parameters for each individual project due to differences in staining protocols, image acquisition and regional differences in staining intensity in subregions of a tissue sample (Roeder et al., 2012). This is a straightforward process in QuPath. Detection parameters may be optimised empirically by adjusting parameters until the detected cells match those observed by the researcher, or can be determined using a more systematic approach, as we employed in this study. As a proof of principle, we undertook a detailed optimisation of the fluorescence intensity thresholds required for accurate detection of cells in all three channels. To expedite this process, we designed a custom script to enable rapid testing of multiple fluorescent intensity thresholds, without having to manually alter them. This script is easily adaptable to accommodate other cell detection parameters available in QuPath. The data we obtained using this approach (Figure 4) enabled us to determine that the fluorescent intensity thresholds for accurate cell detection were different in the various sub-regions we analysed in our coronal mouse brain slices. This conclusion was reached by comparing the automated counts at different thresholds to manual counts, with the assumption that our manual counts represent ground truth.

We have not yet determined why different sub-regions required different intensity thresholds for accurate cell detection in our coronal mouse brain sections. This could be a biological feature of the tissue. For instance, microglia in the thalamus may express a different level of CX_3_CR1-GFP than cells in the cortex, therefore requiring a different intensity threshold for accurate quantification. Alternatively, it could be a technical artefact. For example, the edges of the tissue often have higher background fluorescence than regions in the middle of the tissue, which may enhance the fluorescence across that entire region, thereby requiring a higher intensity threshold to detect cells without also detecting background fluorescence. Whatever the cause, the finding that different sub-regions of interest required different cell detection parameters highlights the importance of optimising detection parameters in each experiment, especially when attempting to compare cell numbers from region to region.

The classification of cells with two separate fluorophores in a single section raises the possibility some cells may be ‘dual-classified’ – that is: a single cell may be detected as expressing both markers (Figure 3 E-F). Whether these are truly cells expressing both markers, or whether this is merely an artefact of the imaging and analysis process, will depend on the markers in question. In our study we did not expect DsRed and GFP to co-localise as NG2 and CX_3_CR1, considered markers of pericytes and microglia in the brain respectively, have not, to the best of our knowledge, been reported to co-localise in the same cells in the healthy adult brain. There is one report of NG2 positive OPCs being engulfed by CX_3_CR1-GFP positive amoeboid microglia in the *corpus callosum* of developing mouse brains. Furthermore, there are reports suggesting pericytes may differentiate into microglia in disease states and thereby begin expressing microglial markers (Ozen et al., 2014; Sakuma et al., 2016). Conversely, others have reported expression of NG2 in microglia (Huang et al., 2020; Zhu et al., 2016). In our tissue however, a visual inspection of the rare dual-classified DsRed and GFP positive cells revealed these were individual DsRed and GFP positive cells with nuclei that were in close proximity (Figure 3 E-F), preventing the automated cell detection from separating the nuclei and consequently classifying them as the same cell. Although these dual classified cells were a rare occurrence, the inability of QuPath (and other automated cell detection programs) to accurately segregate close nuclei remains one of the limitations of automated cell counting. This limitation is best overcome by manual cell counting approaches, such as stereology.

We quantified microglia and pericytes in several brain regions, observing some significant differences between regions. Microglia are thought to account for ~10% of the total number cells in the human brain and ~5-10% in the mouse brain, although these numbers vary across brain regions (Lawson et al., 1990; von Bartheld et al., 2016). These numbers are consistent with our study where we found 4.51% of total cells were microglia, with the highest prevalence of GFP-positive microglia in the cortex. The precise percentage of cells that are microglia in mouse brains has rarely been quantified. Recently, Dos Santas et al., reported that microglia represent ~6% of all cells in the mammalian cerebral cortical gray matter, after pooling data from 30 species (Dos Santos et al., 2020). For mice specifically, one previous study reported F4/80+ microglia accounted for ~5-12% of the total number of cells in the mouse brain, depending on the region analysed (Lawson et al., 1990). In particular, Lawson et al., found ~5% of cells in the cerebral cortex were F4/80+ microglia, which compares favourably to our data from the cortex (5.54% of total cell detections). The small differences between our studies and others may be accounted for by the precise anatomical regions analysed, our use of the CX_3_CR1 promoter to drive GFP expression in microglia, differences in tissue processing, quantification methodologies and the strain of mouse we used mice (C57/BL6).

The precise percentage of brain cells that are pericytes is more difficult to compare as the quantification of pericytes has not traditionally been included in most brain cell counting studies (von Bartheld et al., 2016). It is estimated that endothelial cells, which form blood vessels and upon which pericytes reside, account for ~30% of non-neuronal cells in the brain (von Bartheld et al., 2016). Considering pericytes provide extensive coverage of endothelial cells (Berthiaume et al., 2018), and that there is an approximate ratio of one pericyte for every three endothelial cells in the brain (Pardridge, 1999), this equates to approximately 5% of all cells in whole brain sections that are possibly pericytes. This is consistent with our study where 3.71% of all cells were detected as pericytes, albeit with the caveat that a small proportion of the NG2 positive cells in our tissue are possibly OPCs (i.e. NG2 glia). In our study, the thalamus was found to have over twice the proportion of DsRed-positive cells compared to the other brain regions assessed. This may be due to the mouse thalamus having an increased vascular volume compared to other brain regions (Xiong et al., 2017), which could reflect the high amount of information that gets transmitted through the thalamus into other brain regions (Sherman and Guillery, 2002). Therefore, pericytes may play an active role in providing energy supply to this important brain region.

The relative number and spatial distribution of both microglia and pericytes can be drastically altered in disease states. Pericyte dysfunction and death are implicated in the pathogenesis of various brain diseases including stroke (Hall et al., 2014) and Alzheimer’s disease (Nortley et al., 2019), while microglia can readily migrate and alter their morphology and function and in disease states (Bachiller et al., 2018). Therefore, the development of rapid and reliable tools like QuPath to quantify alterations in these cell populations will enhance our understanding of these cells in both health and disease.

Although our manual and automated cell counts were highly correlated, cell detection with QuPath is not yet perfect, as illustrated by the examples in Figure 3. It is not yet clear whether the error rate in automated cell counting is significantly different from the error rate inherent in manual counting. Only formal comparisons across multiple operators and multiple automated settings in software like QuPath can answer the question of whether rapid automated cell counting will ever replace current gold standard manual cell counting techniques (ie: stereological cell counting).

Here, we have followed a simple cell classification workflow in order to demonstrate the potential of QuPath as a research tool. Moving forward, QuPath offers a number of tools which could improve cell quantification further. We largely avoided inter- and intra-sample variability in fluorescence quality through our use of endogenous fluorophore expression, so simple fluorescent intensity thresholding was accurate enough for our analysis. More refined results may be possible however using the machine learning algorithms that are built into QuPath. These trainable algorithms are likely to be particularly useful for creating classifiers capable of identifying cells positive for specific markers in tissue with varying degrees of immunofluorescent staining/imaging quality in different biological samples, for example differing levels of background artefacts such as age-related lipofuscin autofluorescence. In addition, QuPath interacts with ImageJ (among other packages), opening increased possibilities for analysis outside the simple methods shown here. Even within our workflow, QuPath generates more data than we have presented. For each detected cell, numerous other parameters are automatically measured including nuclear size and shape, and XY coordinates that can be used for further spatial analysis. Some of these analyses are already built into QuPath and others could be scripted as required or the data exported for analysis elsewhere. Further development of this methodology could include automating the analysis of tissue sections that have been imaged across multiple planes.

In conclusion, QuPath offers a user-friendly solution to whole-slide image analysis which will decrease reliance on down-sampling and region-of-interest analysis. Our workflow illustrates the quick and efficient detection of both pericytes and microglia in the mouse brain and we were able to detect significant differences in microglial and pericyte cell numbers in different brain regions. The workflows we employed, and other functions within QuPath, make this a reliable automated image analysis tool for cell counting in fluorescently-labelled tissue that could lead to important new discoveries in both health and disease.

## Acknowledgements

The authors would like to thank Peter Bankhead and other contributors from the Image.sc forum for their assistance with QuPath script development and Jenna Ziebell for providing the CX_3_CR1^GFP/GFP^ transgenic mice.

---

**Figure 2-1.**
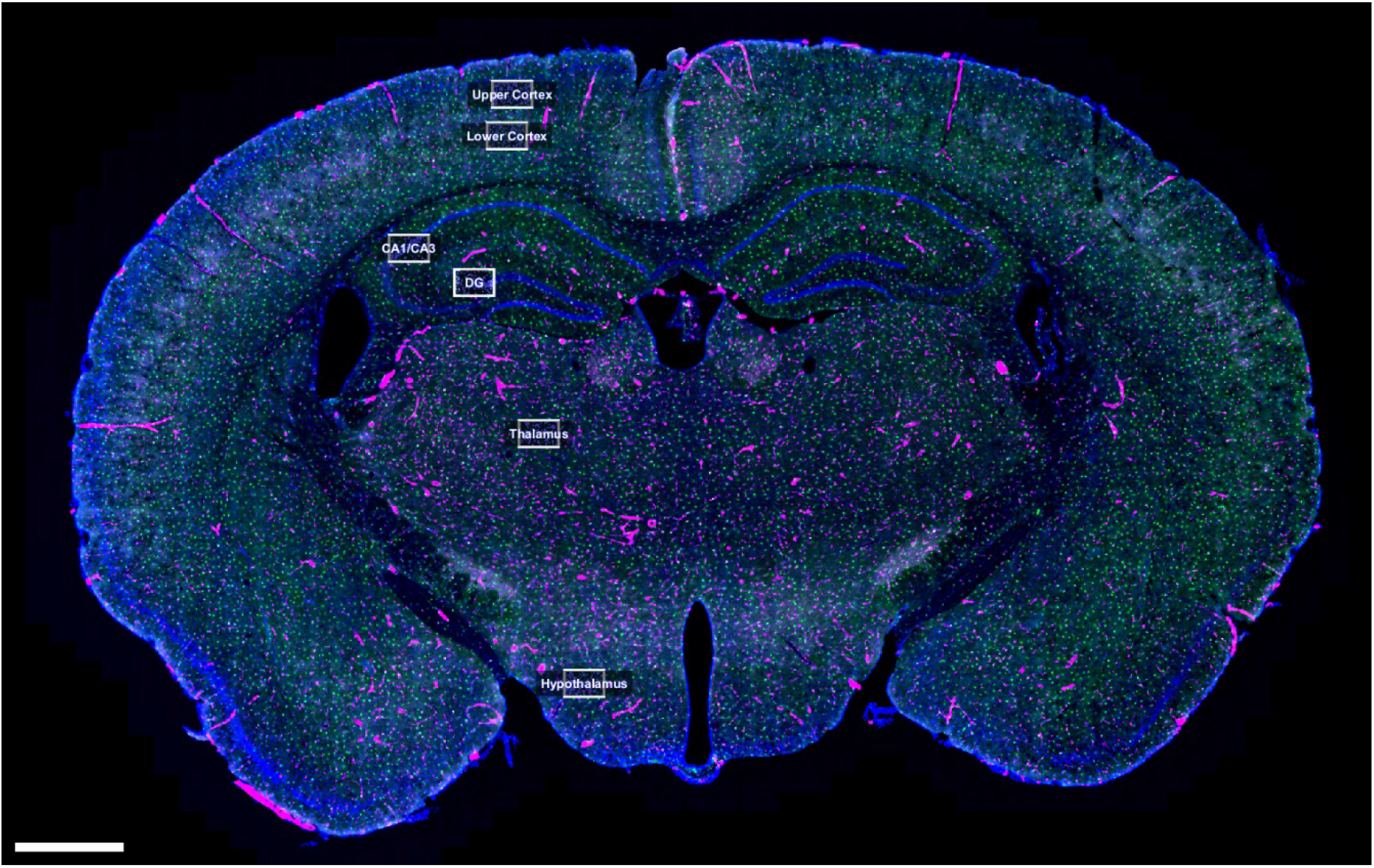
Example of optimisation annotations. 300 x 200μm regions of interest were placed in the upper cortex (layers 1-3), lower cortex (layers 4-6), hippocampus (including dentate gyrus), hippocampus (including CA1/CA3 boundary), thalamus, and hypothalamus of each brain section for the purposes of manually counting cells. Scale bar = 800μm.

**Figure 4-1.**
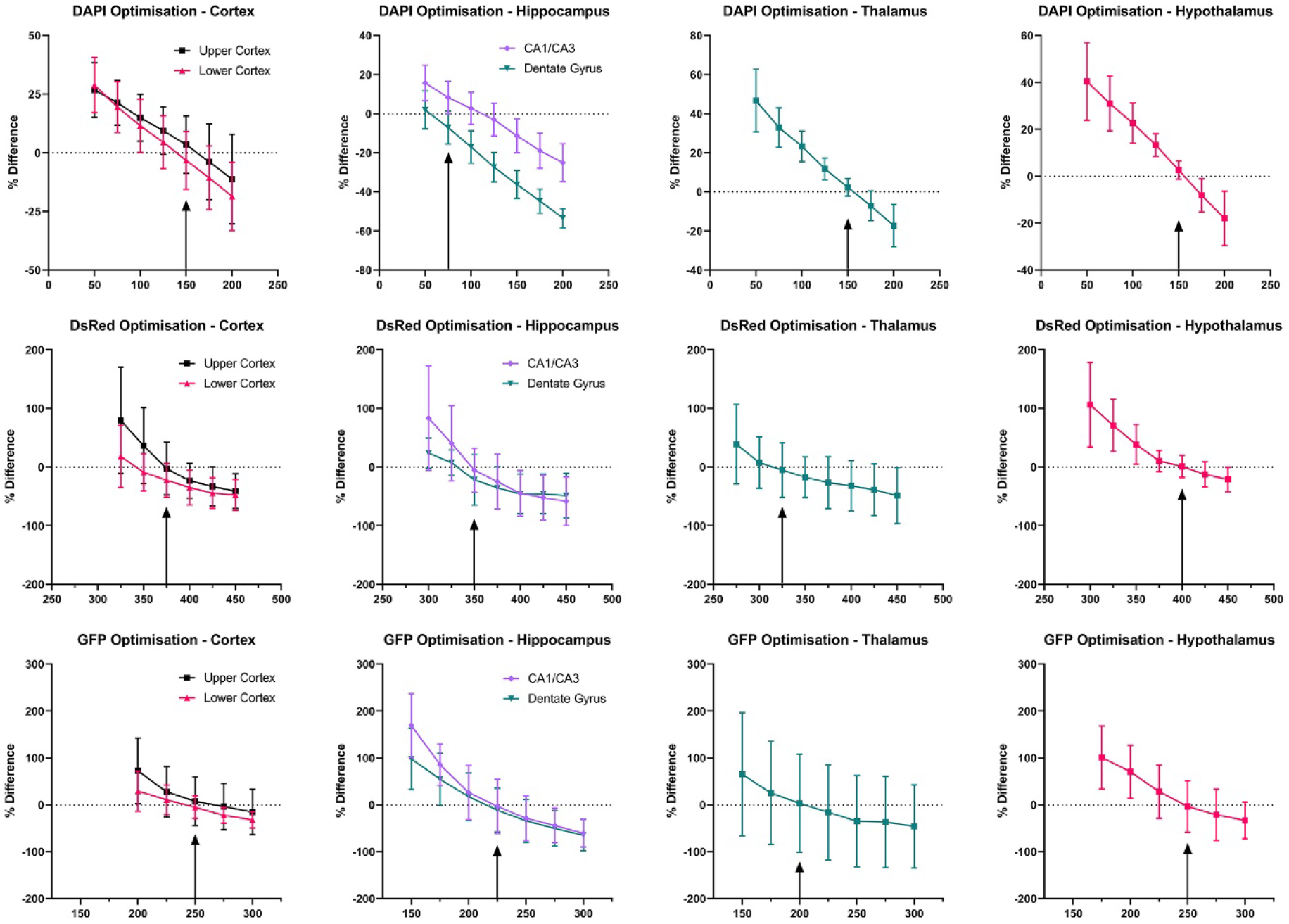
Detailed optimisation of cell detection and classification thresholds. Counts generated by QuPath’s Cell Detection/Positive Cell Detection algorithm were compared to manual cell counts to generate a % difference (dotted line at 0%) with a range of intensity thresholds across six brain regions for DAPI, DsRed, GFP (n=8, mean ± standard deviation). Data for lower thresholds with large standard deviations have been excluded in order to clearly determine the optimum threshold for each region and channel (indicated with an arrow).

**Figure 4-2.**
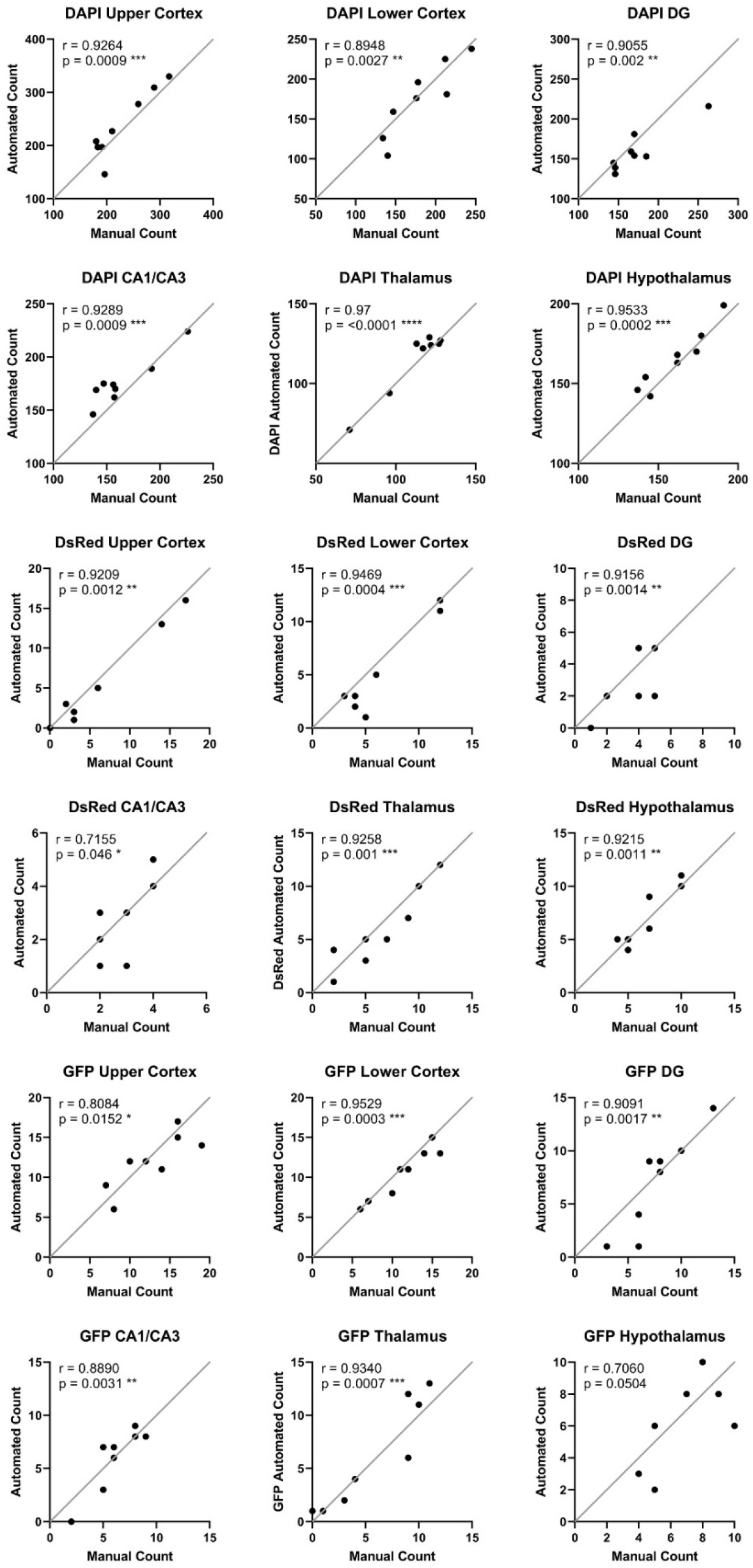
Correlation of manual counts to automated counts at final optimised thresholds. For each optimised threshold, the Pearson’s correlation coefficient (r) between cells counted manually and by QuPath was calculated.

**Figure 5-1.**
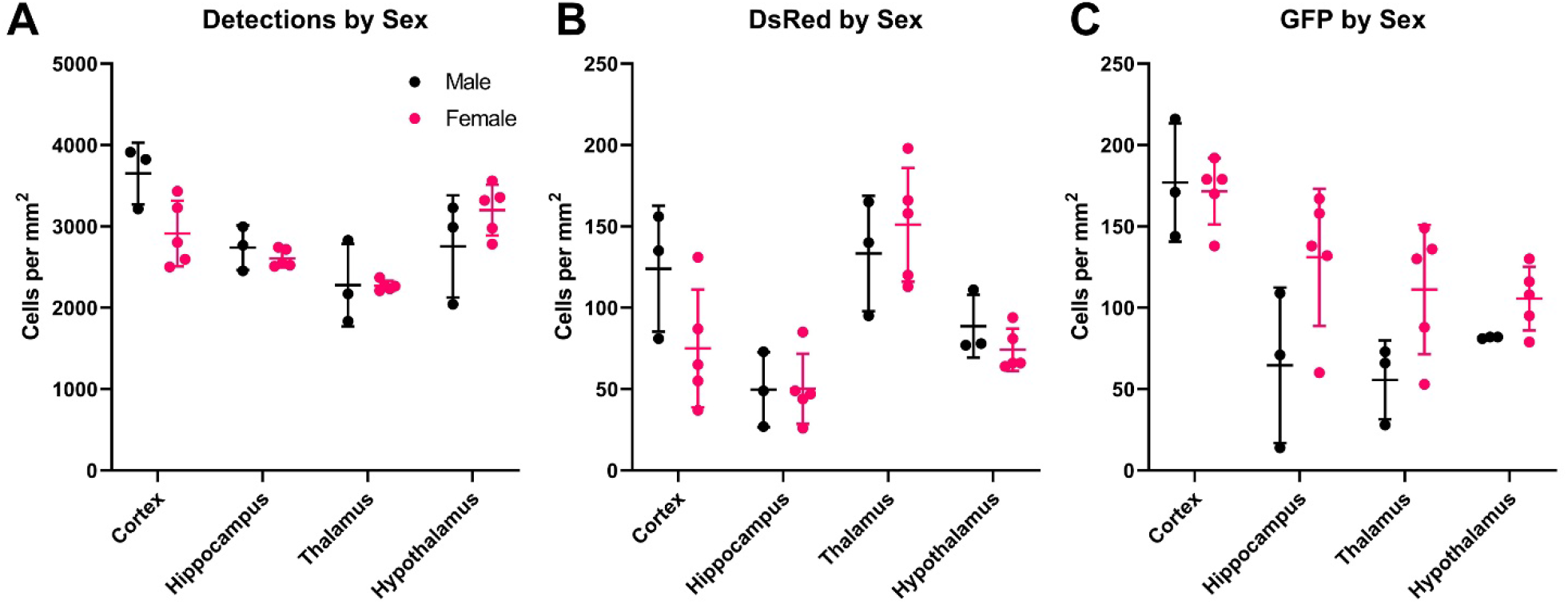
Detection and classification of cells by sex. A) Total cells; B) DsRed-positive cells; and C) GFP-positive cells detected per mm^2^ tissue area. No effect of sex was found by two-way ANOVA.

## References

Al-Janabi, S., Huisman, A. and Van Diest, P.J., 2012. Digital pathology: current status and future perspectives. Histopathology. 61, 1–9.

Armulik, A., Genove, G., Mae, M., Nisancioglu, M.H., Wallgard, E., Niaudet, C., He, L., Norlin, J., Lindblom, P., Strittmatter, K., Johansson, B.R. and Betsholtz, C., 2010. Pericytes regulate the blood-brain barrier. Nature. 468, 557–61.

Bachiller, S., Jiménez-Ferrer, I., Paulus, A., Yang, Y., Swanberg, M., Deierborg, T. and Boza-Serrano, A., 2018. Microglia in Neurological Diseases: A Road Map to Brain-Disease Dependent-Inflammatory Response. Frontiers in Cellular Neuroscience. 12.

Bankhead, P., 2020. Multiplexed analysis — QuPath 0.2.3 documentation.

Bankhead, P., Fernandez, J.A., McArt, D.G., Boyle, D.P., Li, G., Loughrey, M.B., Irwin, G.W., Harkin, D.P., James, J.A., McQuaid, S., Salto-Tellez, M. and Hamilton, P.W., 2018. Integrated tumor identification and automated scoring minimizes pathologist involvement and provides new insights to key biomarkers in breast cancer. Lab Invest. 98, 15–26.

Bankhead, P., Loughrey, M.B., Fernandez, J.A., Dombrowski, Y., McArt, D.G., Dunne, P.D., McQuaid, S., Gray, R.T., Murray, L.J., Coleman, H.G., James, J.A., Salto-Tellez, M. and Hamilton, P.W., 2017. QuPath: Open source software for digital pathology image analysis. Sci Rep. 7, 16878.

Beard, D.J., Brown, L.S. and Sutherland, B.A., 2020. The rise of pericytes in neurovascular research. Journal of Cerebral Blood Flow & Metabolism. 40, 2366–2373.

Berthiaume, A.-A., Grant, R.I., McDowell, K.P., Underly, R.G., Hartmann, D.A., Levy, M., Bhat, N.R. and Shih, A.Y., 2018. Dynamic Remodeling of Pericytes In Vivo Maintains Capillary Coverage in the Adult Mouse Brain. Cell Reports. 22, 8–16.

Brown, L.S., Foster, C.G., Courtney, J.-M., King, N.E., Howells, D.W. and Sutherland, B.A., 2019. Pericytes and Neurovascular Function in the Healthy and Diseased Brain. Frontiers in Cellular Neuroscience. 13.

Colonna, M. and Butovsky, O., 2017. Microglia Function in the Central Nervous System During Health and Neurodegeneration. Annual Review of Immunology. 35, 441–468.

Dos Santos, S.E., Medeiros, M., Porfìrio, J., Tavares, W., Pessoa, L., Grinberg, L., Leite, R.E.P., Ferretti-Rebustini, R.E.L., Suemoto, C.K., Filho, W.J., Noctor, S.C., Sherwood, C.C., Kaas, J.H., Manger, P.R. and Herculano-Houzel, S., 2020. Similar Microglial Cell Densities across Brain Structures and Mammalian Species: Implications for Brain Tissue Function. J Neurosci. 40, 4622–4643.

Finney, C.A., Jones, N.M. and Morris, M.J., 2020. A scalable, fully automated approach for regional quantification of immunohistochemical staining of astrocytes in the rat brain. J Neurosci Methods. 348, 108994.

Hall, C.N., Reynell, C., Gesslein, B., Hamilton, N.B., Mishra, A., Sutherland, B.A., O’Farrell, F.M., Buchan, A.M., Lauritzen, M. and Attwell, D., 2014. Capillary pericytes regulate cerebral blood flow in health and disease. Nature. 508, 55–60.

Huang, W., Bai, X., Meyer, E. and Scheller, A., 2020. Acute brain injuries trigger microglia as an additional source of the proteoglycan NG2. Acta Neuropathol Commun. 8, 146.

Humphries, M.P., Hynes, S., Bingham, V., Cougot, D., James, J., Patel-Socha, F., Parkes, E.E., Blayney, J.K., O’Rorke, M.A., Irwin, G.W., McArt, D.G., Kennedy, R.D., Mullan, P.B., McQuaid, S., Salto-Tellez, M. and Buckley, N.E., 2018. Automated Tumour Recognition and Digital Pathology Scoring Unravels New Role for PD-L1 in Predicting Good Outcome in ER-/HER2+ Breast Cancer. J Oncol. 2018, 2937012.

Lawson, L.J., Perry, V.H., Dri, P. and Gordon, S., 1990. Heterogeneity in the distribution and morphology of microglia in the normal adult mouse brain. Neuroscience. 39, 151–70.

Ledys, F., Klopfenstein, Q., Truntzer, C., Arnould, L., Vincent, J., Bengrine, L., Remark, R., Boidot, R., Ladoire, S., Ghiringhelli, F. and Derangere, V., 2018. RAS status and neoadjuvant chemotherapy impact CD8+ cells and tumor HLA class I expression in liver metastatic colorectal cancer. J Immunother Cancer. 6, 123.

Lein, E.S., Hawrylycz, M.J., Ao, N., Ayres, M., Bensinger, A., Bernard, A., Boe, A.F., Boguski, M.S., Brockway, K.S., Byrnes, E.J., Chen, L., Chen, L., Chen, T.M., Chin, M.C., Chong, J., Crook, B.E., Czaplinska, A., Dang, C.N., Datta, S., Dee, N.R., Desaki, A.L., Desta, T., Diep, E., Dolbeare, T.A., Donelan, M.J., Dong, H.W., Dougherty, J.G., Duncan, B.J., Ebbert, A.J., Eichele, G., Estin, L.K., Faber, C., Facer, B.A., Fields, R., Fischer, S.R., Fliss, T.P., Frensley, C., Gates, S.N., Glattfelder, K.J., Halverson, K.R., Hart, M.R., Hohmann, J.G., Howell, M.P., Jeung, D.P., Johnson, R.A., Karr, P.T., Kawal, R., Kidney, J.M., Knapik, R.H., Kuan, C.L., Lake, J.H., Laramee, A.R., Larsen, K.D., Lau, C., Lemon, T.A., Liang, A.J., Liu, Y., Luong, L.T., Michaels, J., Morgan, J.J., Morgan, R.J., Mortrud, M.T., Mosqueda, N.F., Ng, L.L., Ng, R., Orta, G.J., Overly, C.C., Pak, T.H., Parry, S.E., Pathak, S.D., Pearson, O.C., Puchalski, R.B., Riley, Z.L., Rockett, H.R., Rowland, S.A., Royall, J.J., Ruiz, M.J., Sarno, N.R., Schaffnit, K., Shapovalova, N.V., Sivisay, T., Slaughterbeck, C.R., Smith, S.C., Smith, K.A., Smith, B.I., Sodt, A.J., Stewart, N.N., Stumpf, K.R., Sunkin, S.M., Sutram, M., Tam, A., Teemer, C.D., Thaller, C., Thompson, C.L., Varnam, L.R., Visel, A., Whitlock, R.M., Wohnoutka, P.E., Wolkey, C.K., Wong, V.Y. et al., 2007. Genome-wide atlas of gene expression in the adult mouse brain. Nature. 445, 168–76.

Loughrey, M.B., Bankhead, P., Coleman, H.G., Hagan, R.S., Craig, S., McCorry, A.M.B., Gray, R.T., McQuaid, S., Dunne, P.D., Hamilton, P.W., James, J.A. and Salto-Tellez, M., 2018. Validation of the systematic scoring of immunohistochemically stained tumour tissue microarrays using QuPath digital image analysis. Histopathology. 73, 327–338.

Morris, G.P., Clark, I.A., Zinn, R. and Vissel, B., 2013. Microglia: A new frontier for synaptic plasticity, learning and memory, and neurodegenerative disease research. Neurobiology of Learning and Memory. 105, 40–53.

Nortley, R., Korte, N., Izquierdo, P., Hirunpattarasilp, C., Mishra, A., Jaunmuktane, Z., Kyrargyri, V., Pfeiffer, T., Khennouf, L., Madry, C., Gong, H., Richard-Loendt, A., Huang, W., Saito, T., Saido, T.C., Brandner, S., Sethi, H. and Attwell, D., 2019. Amyloid β oligomers constrict human capillaries in Alzheimer’s disease via signaling to pericytes. Science. 365, eaav9518.

Ozen, I., Deierborg, T., Miharada, K., Padel, T., Englund, E., Genove, G. and Paul, G., 2014. Brain pericytes acquire a microglial phenotype after stroke. Acta Neuropathol. 128, 381–96.

Pardridge, W.M., 1999. Blood-brain barrier biology and methodology. J Neurovirol. 5, 556–69.

Roeder, A.H., Cunha, A., Burl, M.C. and Meyerowitz, E.M., 2012. A computational image analysis glossary for biologists. Development. 139, 3071–80.

Sakuma, R., Kawahara, M., Nakano-Doi, A., Takahashi, A., Tanaka, Y., Narita, A., Kuwahara-Otani, S., Hayakawa, T., Yagi, H., Matsuyama, T. and Nakagomi, T., 2016. Brain pericytes serve as microglia-generating multipotent vascular stem cells following ischemic stroke. J Neuroinflammation. 13, 57.

Sherman, S.M. and Guillery, R.W., 2002. The role of the thalamus in the flow of information to the cortex. Philosophical transactions of the Royal Society of London. Series B, Biological sciences. 357, 1695–1708.

Sweeney, M.D., Ayyadurai, S. and Zlokovic, B.V., 2016. Pericytes of the neurovascular unit: key functions and signaling pathways. Nat Neurosci. 19, 771–83.

von Bartheld, C.S., Bahney, J. and Herculano-Houzel, S., 2016. The search for true numbers of neurons and glial cells in the human brain: A review of 150 years of cell counting. The Journal of comparative neurology. 524, 3865–3895.

Xiong, B., Li, A., Lou, Y., Chen, S., Long, B., Peng, J., Yang, Z., Xu, T., Yang, X., Li, X., Jiang, T., Luo, Q. and Gong, H., 2017. Precise Cerebral Vascular Atlas in Stereotaxic Coordinates of Whole Mouse Brain. Front Neuroanat. 11, 128.

Zhu, L., Su, Q., Jie, X., Liu, A., Wang, H., He, B. and Jiang, H., 2016. NG2 expression in microglial cells affects the expression of neurotrophic and proinflammatory factors by regulating FAK phosphorylation. Sci Rep. 6, 27983.

